# In vivo assessment of mechanisms underlying the neurovascular basis of postictal amnesia

**DOI:** 10.1101/2020.01.30.926717

**Authors:** Jordan S. Farrell, Roberto Colangeli, Barna Dudok, Marshal D. Wolff, Sarah L. Nguyen, Jesse Jackson, Clayton T. Dickson, Ivan Soltesz, G. Campbell Teskey

## Abstract

Long-lasting confusion and memory difficulties during the postictal state remain a major unmet problem in epilepsy that lacks pathophysiological explanation and treatment. We previously identified that long-lasting periods of severe postictal hypoperfusion/hypoxia, not seizures *per se*, are associated with memory impairment after temporal lobe seizures. While this observation suggests a key pathophysiological role for insufficient energy delivery, it is unclear how the networks that underlie episodic memory respond to vascular constraints that ultimately give rise to amnesia. Here, we focused on cellular/network level analyses in the CA1 of hippocampus *in vivo* to determine if neural activity, network oscillations, synaptic transmission, and/or synaptic plasticity are impaired following kindled seizures. Importantly, the induction of severe postictal hypoperfusion/hypoxia was prevented in animals treated by a COX-2 inhibitor, which experimentally separated seizures from their vascular consequences. We observed complete activation of CA1 pyramidal neurons during brief seizures, followed by a short period of reduced activity and flattening of the local field potential that resolved within minutes. During the postictal state, constituting tens of minutes to hours, we observed no changes in neural activity, network oscillations, and synaptic transmission. However, long-term potentiation of the temporoammonic pathway to CA1 was impaired in the postictal period, but only when severe local hypoxia occurred. Lastly, we tested the ability of rats to perform object-context discrimination, which has been proposed to require temporoammonic input to differentiate between sensory experience and the stored representation of the expected object-context pairing. Deficits in this task following seizures were reversed by COX-2 inhibition, which prevented severe postictal hypoxia. These results support a key role for hypoperfusion/hypoxia in postictal memory impairments and identify that many aspects of hippocampal network function are resilient during severe hypoxia except for long-term synaptic plasticity.

## Introduction

The postictal state is marked by brain region-specific dysfunction, can last up to several hours, and be a significant cause of morbidity in epilepsy^1-3^. While the principal goal of epilepsy therapy is to completely prevent seizures, this is not attained in 30-40% of patients, including up to 75% of persons with lesional temporal lobe epilepsy^4^. Modern antiseizure medications have not reduced the proportion of drug refractory epilepsy^5^ and periods of postictal confusion and memory difficulties are a common source for lowered quality of life. A major unmet need not covered in modern epilepsy therapies is the prevention of postictal impairments, which long outlast seizures and are a major hurdle impeding the return to daily life. Thus, the postictal state represents an opportunity for fundamentally new treatments.

Defining the postictal state has been difficult because of the lack of a pathophysiological biomarker. Postictal EEG often detects a suppression of activity beginning immediately after seizure termination, but this lasts less than a few minutes^6^. Critically, postictal amnesia and other cognitive/behavioral impairments occur on a vastly longer time-scale of tens of minutes to hours when clear EEG abnormalities are not observed^7^. While the pathophysiological underpinnings of postictal impairments remain elusive, recent data highlight a potential causal role for severe and local postictal hypoperfusion/hypoxia that occurs in rodents and people^8-10^. Therefore, postictal vasoconstriction-induced hypoperfusion/hypoxia could provide an objective pathophysiological biomarker for defining the postictal period^11^. Indeed, blocking this long-lasting stroke-like event with pharmacological tools (e.g. L-type calcium channel blockers or cyclooxygenase-2, COX-2, inhibitors) prevents the occurrence of postictal behavioral symptoms following focal seizures^8^. Moreover, seizure models without a postictal state do not experience postictal hypoperfusion/hypoxia^12^. While these studies support a central vascular role for postictal behavioral impairment, the underlying mechanisms are not fully understood and must be resolved to better understand the consequences of severe postictal hypoperfusion/hypoxia on synaptic plasticity and memory formation.

Here, we used the COX inhibitor, acetaminophen, as a tool to block hypoperfusion/hypoxia following brief non-convulsive hippocampal seizures to understand the effect of severe hypoxia (pO_2_ < 10mmHg) on hippocampal physiology and behavior. Experiments were performed *in vivo* to maintain an intact neurovascular unit and were designed to address which aspects of hippocampal network function, including CA1 pyramidal neuron activity, local extracellular oscillations, synaptic function, and long-term potentiation (LTP), are impaired following seizures. Only LTP, but not the other metrics of network function, was impaired postictally in a hypoxia-dependent manner. Like LTP impairment, postictal amnesia was prevented by acetaminophen pre-treatment. These results provide insight into how seizures, but more importantly the resulting stroke-like event, lead to long-lasting postictal memory impairments and form the basis for a unique class of epilepsy therapies that target the secondary consequences of seizures when seizure control is not achieved.

## Results

### Kindled seizures lead to prolonged vasoconstriction that does not suppress neuronal activity

We first tested the hypothesis that severe hypoperfusion/hypoxia in the postictal state suppresses neuronal activity, which is expected to occur under conditions of poor neurovascular coupling^13^. Since COX inhibitors have been previously shown to prevent postictal hypoperfusion/hypoxia^8^, we used acetaminophen as a tool to inhibit COX-2 and result in seizures with hypoperfusion/hypoxia (vehicle pre-treatment) or without (acetaminophen pre-treatment). Neuronal activity was measured by 2-photon imaging of somatic calcium activity with GCaMP6f^14^ simultaneously in hundreds of CA1 pyramidal neurons from awake, head-fixed mice on a treadmill (Fig. 1A). This approach also enabled the ability to estimate blood vessel diameter (absence of fluorescence) to confirm the occurrence postictal hypoperfusion (Fig. 1A).

**Figure 1:**
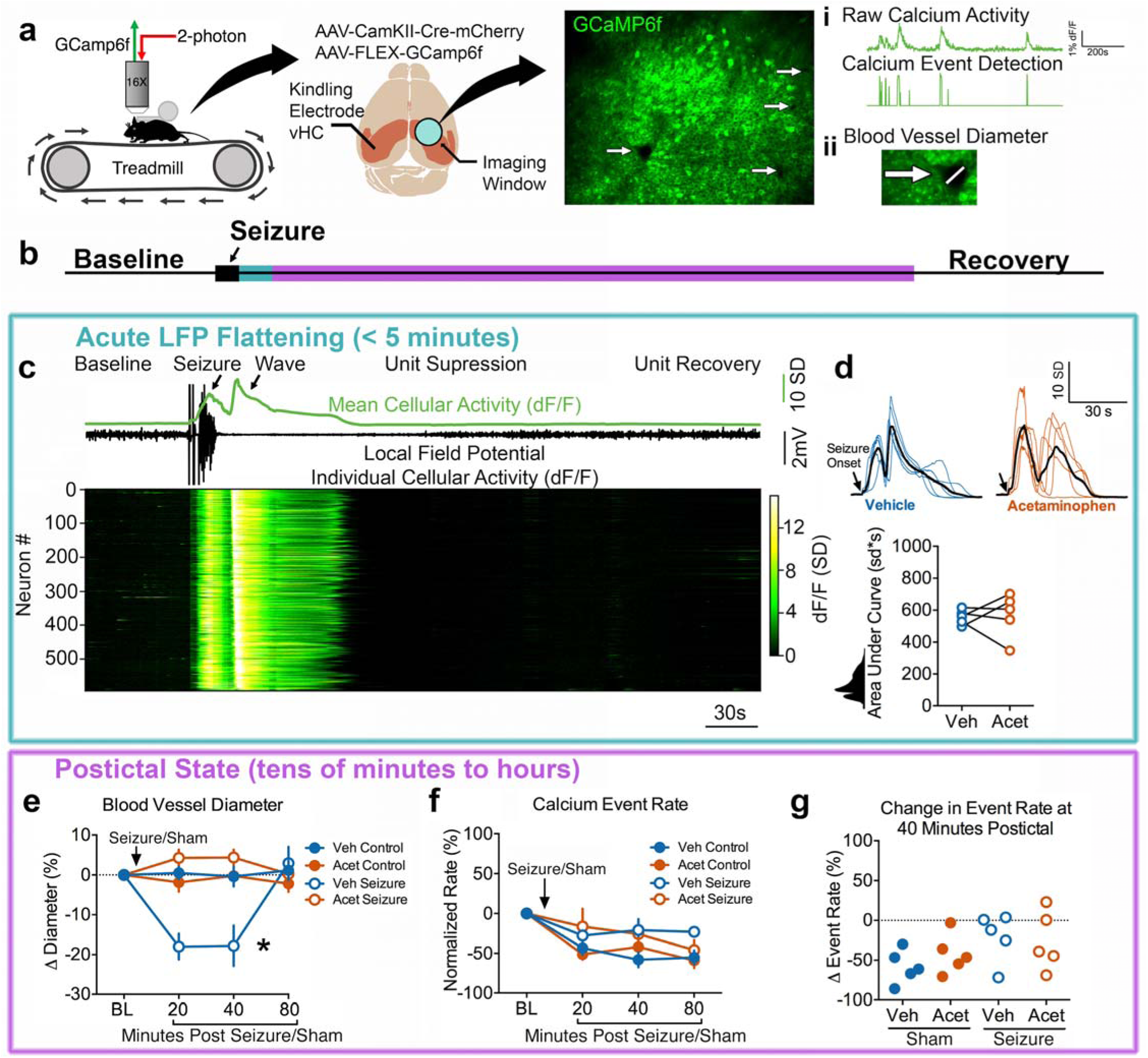
Postictal vasoconstriction does not alter CA1 pyramidal cell calcium activity. (**a**) Experimental set-up. Awake, head-fixed mice moved voluntarily on a linear treadmill while being imaged on a 2-photon microscope. Two viruses were used to achieve GCaMP6f expression in CA1 pyramidal neurons. Seizures were elicited and electrographically recorded in the contralateral ventral hippocampus. Measurements obtained were (i) a change in neuronal calcium activity as a change in GCaMP6f fluorescence, extracted calcium events, and (ii) blood vessel diameter as seen by an absence of GCaMP6f expression. (**b**) Relative timeline associated with the dependent measurements in the following panels corresponding to acute LFP flattening (blue; **c** and **d**) and the longer postictal state (purple; **e**-**g**). (**c**) Representative seizure recording from a mouse treated with vehicle. Calcium traces from 594 neurons plotted with the mean change in fluorescence (displayed as z-score; scale shows number of standard deviations (SD)) and LFP. Neurons are arranged by the timing of wave onset. Seizures resulted in recruitment of all neurons, followed by an intense prolonged calcium wave, suppression of firing, and recovery. (**d**) Mean calcium trace from all recorded neurons of each individual mouse (n=5 for each) is plotted and aligned at seizure onset. Colored traces are from individual mice pre-treated with vehicle (blue; dimethyl sulfoxide) or acetaminophen (orange; 250mg/kg i.p.) with each group mean overlaid in black. Bottom: Quantification. The total area under the curve was not significantly different for pre-treated with acetaminophen or vehicle (t(4)=0.22, p=0.84). (**e**) Change in blood vessel diameter from pre-seizure/sham level baseline (BL). Within subject 2-way ANOVA revealed a significant effect of time (F(3,12)=4.42, p=0.03), condition (F(3,12)=6.38, p=0.008), and an interaction (F(3,12)=12.39, p<0.0001). Tukey post-test demonstrated that vehicle injection with a seizure resulted in a significant decrease in blood vessel relative to vehicle injection without a seizure. Data are mean ± SEM. (**f**) Calcium event rate normalized to baseline. Within subject 2-way ANOVA showed an effect of time (F(3,12)=15.05, p=0.0002), presumably as mice become less aroused over time, but no effect of condition (F(3,12)=3.45, p=0.051) or an interaction (F(9,36)=2.00, p=0.069). Only events during immobility were analyzed here, as movement was not consistently seen across imaging sessions. Data are mean ± SEM. (**g**) At 40 minutes post-seizure (or sham), the percent change from baseline calcium event rate was calculated. No effect of treatment was observed (within subject 1-way ANOVA, F(3)=2.60, p=0.15).

We first determined seizure characteristics, since previous data reports that longer seizures result in more severe hypoperfusion/hypoxia^8^. Seizures were elicited by brief electrical stimulation (kindling: 1ms biphasic pulses <200µA at 60Hz for 1 second to the ventral hippocampus contralateral to the imaging cannula; Fig. 1a) and resulted in short afterdischarges (LFP trace in Fig. 1c) with little to no behavioral manifestation (corresponding to stage 0-1 on the Racine scale^15^). The electrographic seizure duration was not significantly different for vehicle vs. acetaminophen pre-treatment (20.5 ± 4.0 vs. 27.4 ± 4.4 s; t(4)=1.49, p=0.21, paired t-test). We then assessed intracellular calcium activity during seizures to understand the upstream events that lead to COX-2-dependent vasoconstriction and severe hypoxia, since elevated intracellular calcium is necessary to mobilize COX-2 substrates^16-19^. Despite the limited behavioral manifestation induced by these kindled seizures, virtually all recorded neurons in the field of view were profoundly activated during seizures (Fig. 1b; Video 1; 99.6% of units had > 6 sd activity increase). Moreover, the calcium activity after stimulation manifested in two distinct phases: (1) a synchronous increase in calcium activity that coincided with electrographic seizure activity and (2) a spreading wave that was closely associated with flattening of the LFP and is consistent with the occurrence of a spreading depolarization event^20^ (Fig. 1c). The calcium wave, however, resulted in a 3 times greater calcium signal than the seizure itself (3.0 ± 0.46 times greater area under curve, t(4)=6.62, **p=0.003, one-sample t-test, data from vehicle-treated mice; Supplementary Fig. 1), highlighting that most of the calcium activity occurred while the LFP was flat and neuronal activity was suppressed (Fig. 1c). Importantly, the total calcium activity accumulated during both phases were not significantly different with acetaminophen pre-treatment (Fig.1d, though see also Supplementary Fig. 1b,c as individual phases differ). These data also confirm that kindled seizures propagate throughout both hippocampi, as expected^21^, thereby permitting experimentation in the hippocampus contralateral to seizure induction. Given that acetaminophen did not alter seizure duration or the total calcium activity associated with the seizure and spreading depolarization, we next examined acetaminophen’s effects on the much longer timescale of the postictal state (Fig. 1b).

We confirmed that blood vessel diameter was reduced by approximately 15% at 20- and 40-minutes post-seizure and recovered by 80-minutes (Fig. 1e), which is in line with a prior study that rigorously evaluated postictal vasoconstriction *in vivo*^22^. Vasoconstriction was prevented by pre-treatment with acetaminophen (Fig. 1e), illustrating the dissociation of experimental groups (i.e. similar seizure dynamics but either with or without hypoperfusion/hypoxia). We then determined the average calcium event rate across cells (Fig. 1a) and, surprisingly, observed no effect of postictal vasoconstriction (Fig. 1f,g). These results are in line with action potential firing rates recorded from people undergoing epilepsy monitoring, where units are transiently suppressed immediately after a seizure, as seen acutely during the spreading calcium wave in *Figure 1c*, but make a complete recovery within a few minutes^23^. Thus, CA1 pyramidal neuron activity is preserved throughout the extended postictal state, even under conditions of restricted blood flow following seizures.

### Severe postictal hypoxia does not alter local field potential oscillations in CA1

We then assessed potential changes to CA1 network properties more broadly by examining LFP dynamics across the discrete synaptic layers of the CA1 and dentate gyrus (DG) (Fig. 2a). Since LFP oscillations at these laminae reflect the net extracellular effect of the heterogenous and cell type-specific local and long-range synaptic inputs^24-27^, detailed examination of LFP provides a wide and sensitive assay for potential network impairment during postictal hypoperfusion/hypoxia. Simultaneous pO_2_ and LFP recordings were performed under light urethane anesthesia (Fig. 3a), which mimics natural sleep dynamics of cycling between periods of slow oscillation and theta states, accompanied by faster gamma oscillations, and avoids the confound of behaviorally-driven LFP changes associated with awake behavior^28,29^ (Fig. 3b). Despite the occurrence of severe hypoxia that lasted over an hour in the hippocampus (pO_2_ < 10mmHg), no prolonged postictal changes in LFP were observed in both theta and slow oscillation states (Fig. 2b). Furthermore, the amplitude of theta (Fig. 2c) and locations of current sinks and sources (Fig. 2d) did not change across the hippocampal laminae during severe postictal hypoxia (Fig. 2e). Since no postictal LFP changes were seen in awake mice during the previous experiment (Supplementary Fig. 2), anesthesia is unlikely to be a confound.

**Figure 2:**
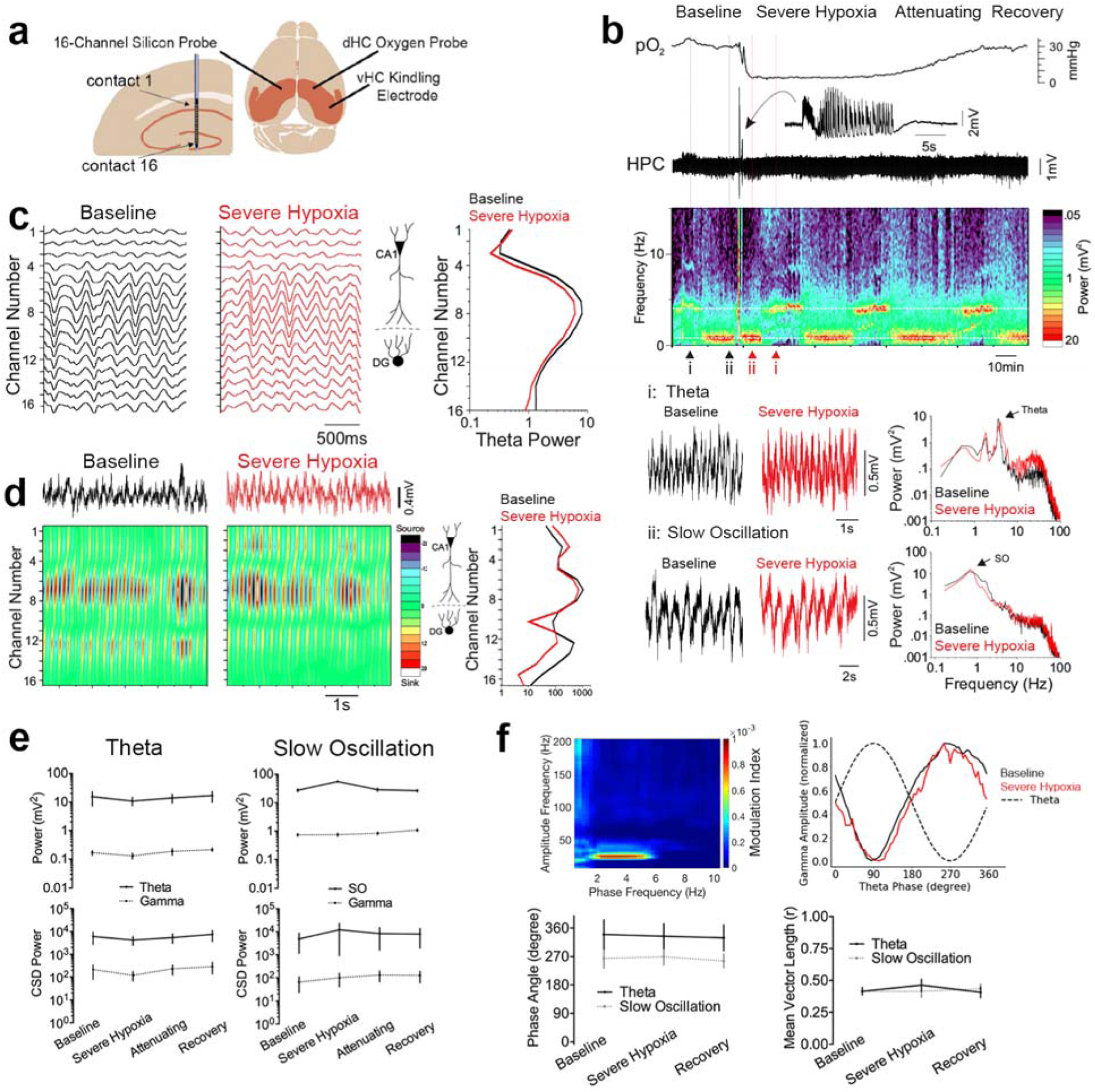
Severe postictal hypoxia does not change CA1 network oscillations. (**a**) Experimental recording paradigm. 16-channel silicon probes were lowered into the CA1 contralateral to the site of seizure initiation and oxygen recording. (**b**) Representative recording of a brief seizure followed by prolonged, severe hypoxia. Raw traces of theta and slow oscillation are displayed according to their position in the spectrogram, which displays cycling between these two states. For these two states, power is plotted against frequency and reveals no amplitude changes across the frequency spectrum. Arrows indicate peak power for each state. (**c**) Filtered theta pre and post seizures (i.e. baseline vs. severe hypoxia) with theta power plotted as a function of depth. The spatial profile of theta power is conserved during severe hypoxic episodes. (**d**) Current source density (CSD) analysis of the theta profile shows overlapping sink and source locations before and during severe hypoxia. (**e**) Quantification of **c** and **d**. 2-way ANOVA revealed no effect of time for theta (n=4) and slow oscillation (n=5) states for raw LFP power (F(3,42)=0.76, p=0.52) and CSD power (F(3,42)=0.39, p=0.76). (**f**) Top left: representative modulation index matrix across an entire recording session showing strong coupling between the low and high frequency LFP components. Top right: representative modulation plot of normalized gamma amplitude (envelope of 20-40Hz filtered signal) plotted against the phase of the theta wave demonstrating strongest coupling at the trough of theta, which is unchanged during severe postictal hypoxia. Bottom: Summary data from the gamma modulation plots obtained during theta and slow oscillation states. The phase angle (phase at peak gamma amplitude; theta: F(2,3) = 0.44, p=0.66, SO: F(2,4) = 1.71, p=0.24) and vector length (theta: F(2,3) = 0.9, p=0.45, SO: F(2,4) = 0.24, p=0.79) are not significantly different during severe hypoxia. Data are mean ± SEM.

**Figure 3:**
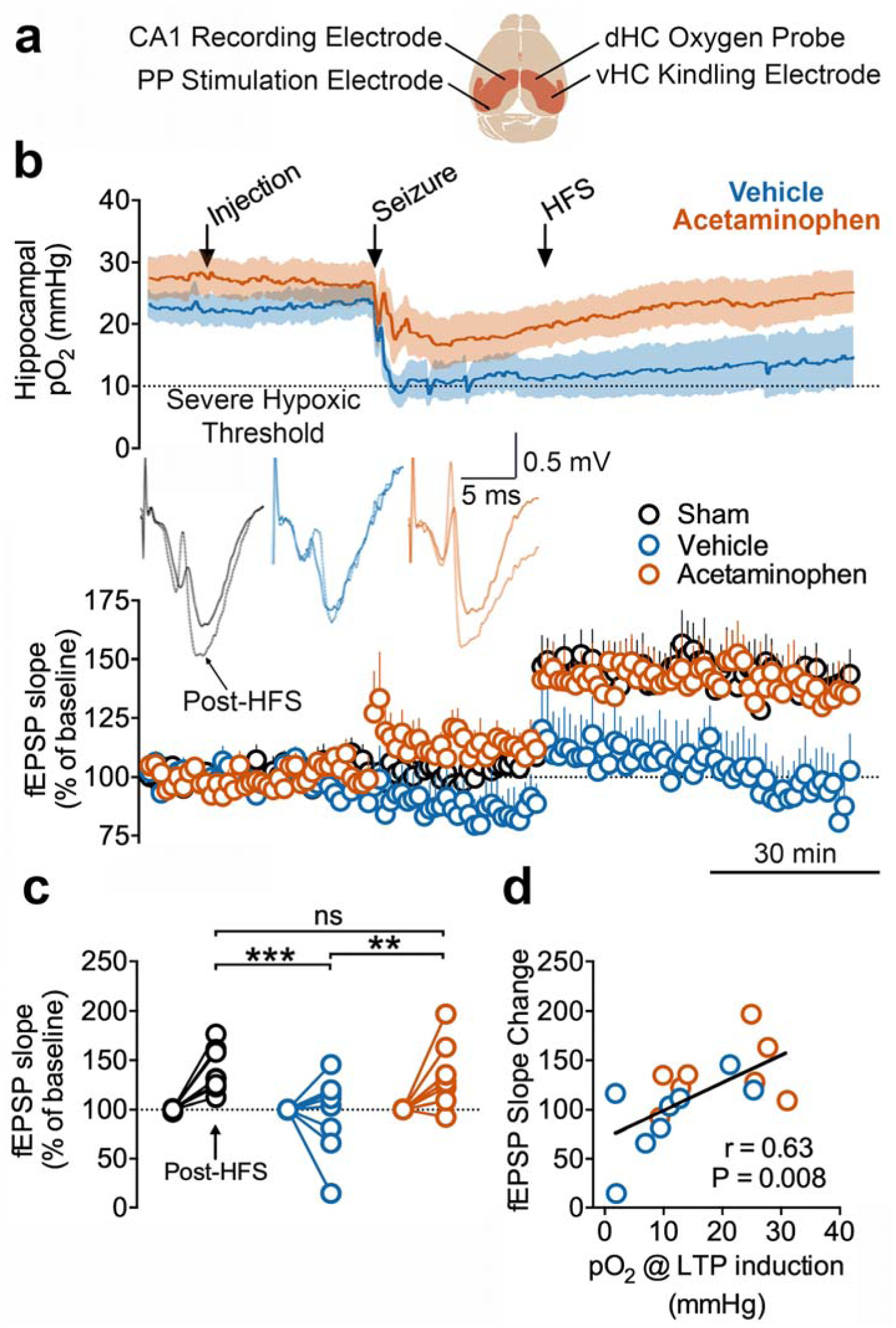
Severe postictal hypoxia prevents LTP. (**a**) Recording paradigm. The right hemisphere was used for seizure induction and oxygen monitoring while the left hemisphere was used from electrophysiology of the PP-CA1 synapse. Responses were collected under urethane anesthesia from an acute electrode aim at the stratum lacunosum moleculare of CA1. fEPSPs were evoked every 20 s and 3 consecutive responses were then averaged (one per minute). (**b**) Concurrent oxygen and electrophysiological recordings from rats that either went severely hypoxic or not after kindled seizures. Mean fEPSP slope ± SEM for each group is displayed before and after HFS. Inset shows representative evoked potentials from a rat in each group before (pre-injection, 10 min baseline) and after HFS (45-55 min post-HFS). (**c**) Quantification of B. Data from 45-55 minutes post-HFS was compared to the initial baseline (pre-injection). Two-way ANOVA revealed an effect of HFS (F(2,21) = 4.61, p=0.02), group assignment (F(1,21) = 12.3, p=0.02), and an interaction (F(2,21) = 4.61, p=0.02). Tukey multiple comparisons are shown with significance levels (**p<0.01, ***P<0.001). (**d**) The postictal oxygen level at the time of LTP induction had a significant, positive correlation with the change in fEPSP slope.

On a sub-second time-scale, the amplitude of gamma oscillations is modulated with respect to the phase of the ongoing low frequency rhythm. This cross-frequency coupling, which is thought to rely on precisely timed heterogeneous inhibition that coordinates assemblies of excitatory cells at this fast time-scale^30,31^, may be perturbed by severe postictal hypoxia even though the overall power spectrum was unchanged on a longer time-scale. We first addressed this hypothesis by objectively determining the high frequency components that couple most strongly to theta and slow oscillation activity (Fig. 2f, top right). Since we observed strong coupling in the gamma range at 20-40 Hz, we then assessed how the amplitude at this frequency band is modulated within theta and slow oscillation cycles. We observed strong coupling near the trough of each cycle, which did not change during severe postictal hypoxia (Fig. 2f). Thus, the coordinated activity at the microcircuit level that gives rise to cross-frequency coupling is still functional during severe postictal hypoxia. When also considering lack of postictal changes in calcium activity of CA1 pyramidal neurons, these results support the view that the resting network properties of CA1 are insensitive to severe hypoperfusion/hypoxia following seizures.

### Severe postictal hypoxia prevents the induction of LTP

Since no hypoperfusion/hypoxia-dependent changes in resting-state network functions were observed, as seen with cellular activity and LFP, we then postulated that additional demands to the network, beyond that of the resting state, may not be supported during postictal hypoxia. Here, we monitored the synaptic strength of the temporoammonic pathway (entorhinal cortex to CA1 via the perforant pathway), a synapse considered to be an important component for hippocampal computation and memory^32-36^, and tested whether high-frequency stimulation (HFS) could induce LTP following seizures with or without hypoxia (Fig. 3a). As with previous experiments, kindling stimulation was delivered contralaterally so that the temporoammonic pathway under investigation would be affected by seizures and hypoxia, but not directly by kindling stimulation. Indeed, no kindling-induced potentiation^37^ or changes in paired-pulse ratio were observed following kindling stimulation (Supplementary Fig. 3; but see Supplementary Fig. 4 for an outlier with an extremely hypoxic hippocampus and suppressed evoked potentials). The observed lack of change in evoked field potentials, even under conditions of severe hypoxia, is consistent with the unchanged LFP oscillations (Fig. 2, Supplementary Fig. 2), since a seizure-induced alteration in synaptic communication would likely be reflected in the LFP.

We then delivered HFS to determine if the temporoammonic synapse can be strengthened following seizures with and without hypoxia. Long-term potentiation, as measured by a long-lasting increase in fEPSP slope, reliably occurred in sham-treated rats but did not occur in vehicle-treated rats following a seizure (Fig. 3b,c). Importantly, rats pre-treated with acetaminophen did not display severe postictal hypoxia and had synaptic potentiation to a level not significantly different from sham-treated rats (Fig. 3b,c; see Supplementary Fig. 3 for analysis of LTP using post-seizure as baseline). Previous reports have demonstrated an inhibitory effect of COX inhibitors on LTP^38-40^, though other studies demonstrate their ability to rescue of LTP under conditions of pathology^41-43^. Acetaminophen treatment here rescued LTP following seizures, perhaps owing to its ability to prevent postictal hypoxia. This hypothesis is clearly supported by the strong positive correlation of postictal pO_2_ with the change in slope of the fEPSP; strength of potentiation (Fig. 3d). These data support that the expression of severe postictal hypoxia may be necessary for LTP suppression at this synapse.

### Severe postictal hypoxia impairs memory

Lastly, since mechanisms that modulate synaptic strength, such as those involved in LTP, are important for learning and memory, we investigated whether severe postictal hypoxia would impair hippocampal-dependent memory performance. We previously demonstrated the occurrence of postictal object memory deficits that can be reversed by acetaminophen pre-treatment^8^. Here, we assessed associative memory using the object/context mismatch task, which requires a paired representation of objects with an environmental context^44,45^. Moreover, this task requires the detection of differences between an encoded experience and current experience, which is thought to require novel sensory input through the temporoammonic path to CA1 that can be compared to the hippocampal representation from CA3^32,46-49^. In this task, rats explored objects that were paired with distinct environments (e.g. different lighting and behavioral chambers) and revisit one of the environments with one of the previously paired objects substituted for an object of the other context (Fig. 4a). Sham-treated controls demonstrate a clear preference for the object that is unfamiliar to that environmental context (Fig. 4c), indicating learning and memory of the prior object/context pairing. Other rats were administered vehicle or acetaminophen before a seizure to have groups of rats with or without severe hypoxia (Fig. 4b), respectively, while performing the task during the postictal period. Preference for the novel object-context pairing did not occur in vehicle-treated rats but was rescued to control levels in acetaminophen-treated rats (Fig. 4c). These results are congruent with the restoration of temporoammonic LTP by acetaminophen pre-treatment and supports a potential role for hypoperfusion/hypoxia in driving memory impairments by inhibiting synaptic plasticity.

**Figure 4:**
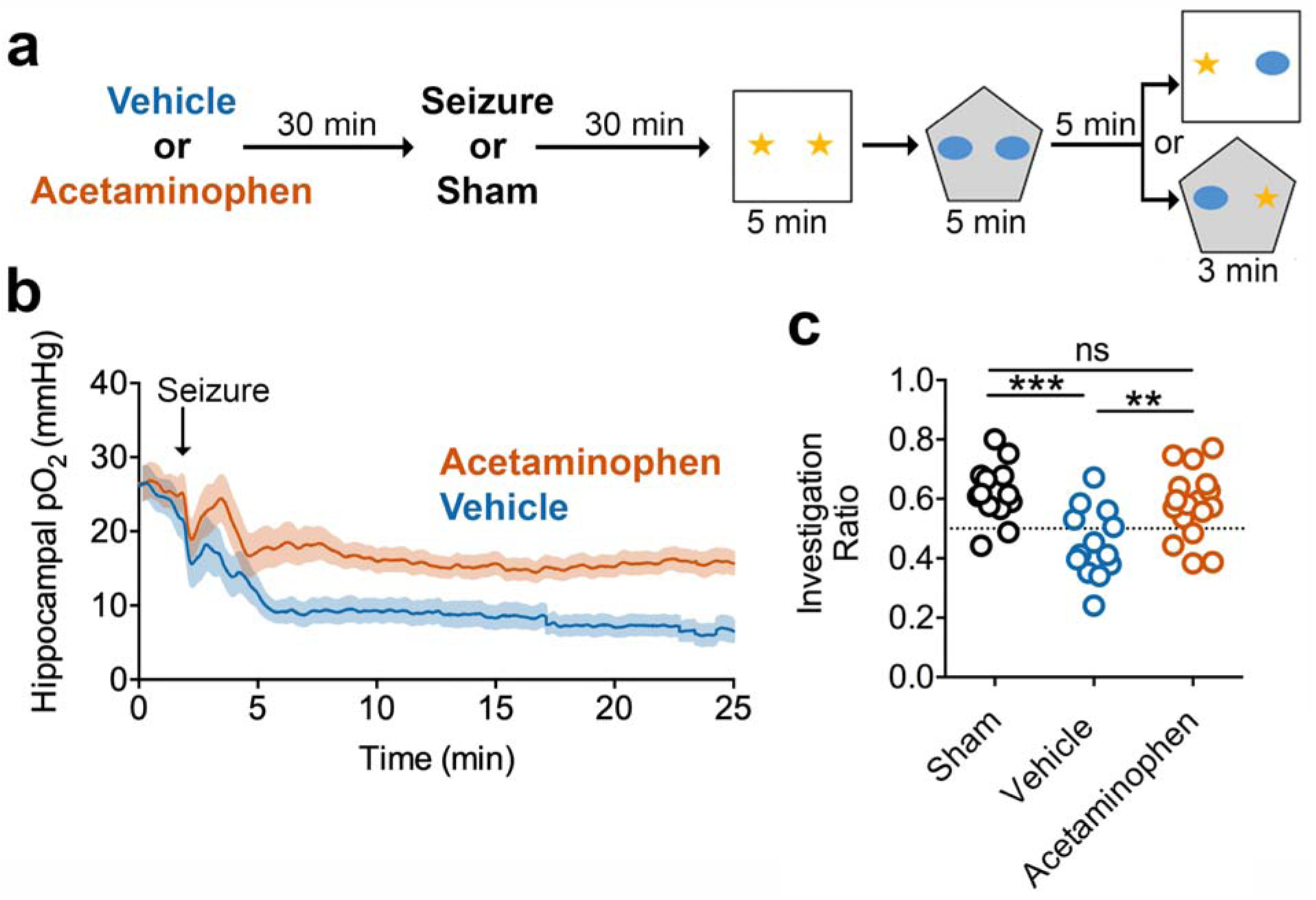
Acetaminophen Prevents Severe Hypoxia and Rescues Postictal Amnesia. (**a**) Experimental paradigm – Object-Context Mismatch. (**b**) Oxygen recordings just prior to the memory task that demonstrate experimental separation of seizure from the resulting hypoxic event by acetaminophen pre-treatment. (**c**) Investigation ratio of the novel object to the familiar object for the paired context. Acetaminophen prevented the seizure-induced memory impairment observed in vehicle-treated rats and performed no differently from sham controls (ANOVA F(2,40)=8.82, follow-up Tukey’s test displayed, ** p<0.01, ***p<0.0001).

## Discussion

Abnormal brain metabolism has long been known to be a contributor to epilepsy pathophysiology^50^. The occurrence of postictal stroke-like events has recently emerged as an important missing link in the pathophysiology of epilepsy^8-11,51-53^ and is a strong candidate mechanism for the occurrence of postictal behavioral impairment. Given the disproportionately high energy requirements of the brain relative to other organs^54^, it is expected that periods of inadequate cerebral blood flow drive neuronal dysfunction, though the impacted subcomponents of neuronal function are less clear. This study examined possible mechanisms that drive postictal amnesia *in vivo* and highlights a potential role for impaired LTP. This research is in line with the observation that synaptic transmission is the most energetically demanding subcellular process in neuronal signaling^55,56^. Since synaptic changes that increase synaptic response amplitude, like LTP, also increase energy consumption^57^, this process puts additional energy burdens on an already demanding process. Our results provide evidence that severe postictal hypoxia does not support synaptic potentiation but is still able to meet the energy requirements to maintain neuronal activity levels, LFP oscillations, and baseline evoked synaptic responses in the CA1. Whether these findings generalize to other hippocampal regions or other brain regions involved in seizures remains to be determined, but these results provide a candidate mechanism for the postictal state that is testable.

The absence of major changes in neuronal activity, baseline synaptic function, and LFP oscillations is perhaps surprising in the context of the extremely low levels of oxygen recorded, since hypoxia is conventionally thought to alter these variables. While anoxia-mimicking conditions are known to profoundly impair synaptic function *in vitro*^57^, those *in vitro* conditions are markedly different from *in vivo* where an approximately 40-50% reduction in blood flow and severe hypoxia locally occur^8-10^, but without complete anoxia. Interestingly, our dataset included one notable outlier with an almost anoxic hippocampus that uniquely demonstrated suppression of synaptic function. This opens the possibility that certain seizures, which drive exceptionally low oxygen levels, have more broad effects beyond LTP prevention and remains to be explored. Our results also differ from whole-body hypoxia exposure, which drives less severe cerebral hypoxia, but is known to drive EEG changes^58,59^. Whereas systemic hypoxia drives a strong homeostatic respiratory and arousal response that affect brain oscillations more globally^60,61^, an important distinction of postictal severe hypoperfusion/hypoxia is its restriction to a few brain regions involved in the seizure^8-10^.

While this study was designed to observe more readily apparent changes in neuronal function, more subtle changes cannot be ruled out. Changes in mean CA1 pyramidal neuronal activity or oscillations across the distinct laminae in this region were not observed, however, it is possible that spike timing in relation to ongoing oscillations is disrupted^25,62,63^. It is also possible that task-specific changes, such as a peak frequency or amplitude shift in theta or gamma frequencies during exploration^26,63^ or neuronal firing corresponding to spatial encoding^65,66^, were selectively impaired and remain to be determined. Changes in these neurophysiological mechanisms could certainly play a role, but the importance of modulating synaptic strength, as identified by this study, cannot be understated.

COX-2 is a crucial enzyme that mediates severe postictal hypoperfusion/hypoxia, whose lipid substrates are mobilized in a calcium-dependent manner^16-19^. Since COX-2 inhibitors given shortly after seizure termination have no effect on postictal hypoxia^8^, the generation of vasoactive prostanoids is likely limited to a very brief time period. An important aspect of this study was the examination of cellular calcium dynamics associated with seizures with high temporal resolution, as intracellular calcium is thought to be a key upstream mediator of postictal hypoperfusion/hypoxia. In addition to the expected increase of neuronal calcium during a seizure, a spreading wave of calcium, consistent with spreading depolarization^20,67^, was consistently observed. This relationship of seizures accompanied by spreading depolarization is often overlooked, as the direct current component of LFP and EEG are rarely recorded but is interestingly noted across many seizure models and clinical epilepsy^68-72^. Since the accumulation of calcium during the wave was three times greater than that of the seizure, spreading depolarization is clearly of considerable relevance for calcium-dependent molecular signaling in epilepsy pathophysiology, such as COX-2^73^ and its crucial role in inducing local vasoconstriction and severe hypoxia that lasts for over an hour^8^. Since spreading depolarization is known to also result in a COX-2-dependent prolonged restriction of blood flow^74,75^, this mechanistic overlap likely exacerbates postictal hypoperfusion/hypoxia when both seizures and spreading depolarization co-occur. These results further clarify the role of COX-2 in coordinating a postictal stroke-like event and highlight its potential as a prime drug target to prevent the secondary effects of seizures.

The rats and mice used in this study were relatively young, but one important future consideration is understanding how these mechanisms operate during aging, since the aged brain is associated with reduced plasticity markers ^76^, cognitive decline, and blunted neurovascular coupling^77^. Interestingly, many of the neurovascular risk factors that are associated with aging are also mechanisms involved in epilepsy^78^. Epilepsy is also associated with a higher incidence in the elderly and the most common risk factor associated with acquiring epilepsy late in life is cerebrovascular disease^79^. Thus, although often overlooked, altered cerebral blood flow regulation is centrally involved in epilepsy and can be both a potential cause and, as outlined here, consequence of seizures.

In summary, these data support a potential synaptic mechanism underlying postictal memory impairment that occurs in a hypoperfusion/hypoxia-dependent manner. One implication from this work is that postictal hypoperfusion/hypoxia could provide an objective pathophysiological biomarker to define the postictal state, as its occurrence is consistently associated with the occurrence of postictal behavioral impairment. This idea is also supported by the recent observation that absence seizures, which are not associated with postictal impairment, are not followed by postictal hypoperfusion/hypoxia^12^. Finally, new antiseizure drugs continue to fail to treat many epilepsies and leave people susceptible to seizures^5^, highlighting that the postictal state still remains a core problem in epilepsy^1-3^. Thus, it is important to have a basic understanding of the secondary effects of seizures and have clinical tools to prevent them. Whether these encouraging results in rodents can be translated to humans is an important next step and is being carried out with clinical trials in Canada (clinicaltrials.gov ID: NCT03949478) and the Netherlands (clinicaltrials.gov ID: NCT04028596).

## Materials and Methods

### Animal Ethics Statement

All experiments performed on rats and mice were approved by Life and Environmental Sciences Animal Care and Health Sciences Animal Care Committees at the University of Calgary and the Administrative Panel on Laboratory Animal Care (APLAC) at Stanford University, respectively, and were performed in accordance with relevant guidelines and regulations. Rats and mice were group-housed in standard cages with unrestricted access to standard chow and water. Rats were single-housed after surgery. All experiments were performed during the light cycle.

### Seizure induction model

To elicit seizures in the rats and mice, we employed electrical kindling targeted at the ventral hippocampus. Note that both rats and mice display postictal hypoxia in a COX-2 dependent manner^8^, which permits experimentation in either species. The chronically implanted electrode in this structure served as both the stimulating and recording electrode, which is facilitated by a switch to direct LFP to an amplifier (Grass Neurodata Acquisition System Model 12C-4-23 for rats, or A-M Systems model 1700 for mice) for recording or allow current to flow from a constant-current stimulator to evoke a seizure (Grass Technologies Model S88 for rats, or A-M Systems model 2100 for mice). Prior to experimentation, all mice and rats underwent a minimum of 5 once daily kindling sessions to evoke seizures by stimulating above the afterdischarge threshold (typically 100-300μA). This was necessary because the first few seizures may be too brief (e.g. under 15 seconds) to result in postictal hypoxia. After a few kindling sessions, seizures are consistently longer and result in more severe postictal hypoxia^8^. Kindling with these few sessions is known to spread bilaterally through both hippocampi^21^, but are associated with low severity, typically stage 0-1 on the Racine scale^15^.

### Awake, head-fixed 2-photon imaging in mice

10 C57BL/6J adult (P90-P120) male mice (Jackson Laboratories, Strain#000664) were injected with 300nL of a 1:1 viral mixture into the CA1 of right dorsal hippocampus (2.2mm posterior, 2.4mm lateral, 1.3mm ventral to bregma). The viral mixture included full titer of AAVDJ-CamKII-mCherry-Cre (Stanford Neuroscience Gene Vector and Virus Core cat#GVVC-AAV-9, a gift generously provided by Karl Deisseroth) and AAV1-Syn-FLEX-GCaMP6f-WPRE-SV40 (Penn Vector Core cat# p2819, now addgene cat# 100833, a gift from The Genetically Encoded Neuronal Indicator and Effector Project (GENIE) & Douglas Kim^14^) to facilitate specific expression in pyramidal neurons. Following one week of recovery a 3mm diameter imaging cannula was implanted as described in Kaifosh et al^80^. Briefly, cannulae were constructed of 3mm glass coverslips (Warner Instruments) mounted to 2 mm long stainless-steel tubing (Tegra Medical). A ∼3mm craniotomy centered at the injection site was performed and the cortex was aspirated. On the contralateral side, a twisted bipolar electrode (Invivo1) with 0.5mm tip separation was implanted into the ventral hippocampus (3.2mm posterior, 2.7mm lateral, 4.0mm ventral to bregma). The implants were secured to the skull with cyanoacrylate glue, dental cement (Lang), and combined with a stainless-steel head-bar for subsequent imaging sessions. Following one to two weeks recovery, all mice underwent 5 once daily kindling sessions (1 second of 60Hz biphasic 1msec square wave pulses) to confirm occurrence of electrographic seizures consistent with ventral hippocampal kindling. We also checked for desirable viral expression with 2-photon microscopy and identified damaged hippocampi that may have occurred during implantation. 5/10 mice moved forward in the study, which began 2-4 weeks after implantation surgery. Reasons for exclusion included misplaced electrodes (2), poor imaging quality due to misplaced virus or damage from surgery (1), or mice removed their head-bar during handling (2).

Mice were acclimated to the imaging set-up for two 10-minute sessions prior to imaging, which consisted of walking/running and immobility on a 1m long belt while head-fixed to minimize the effects of stress during the experiment. Mice served as their own controls and underwent four conditions: (1) vehicle (dimethyl sulfoxide) + control (no seizure), (2) acetaminophen (250mg/kg i.p. from Sigma) + control, (3) vehicle + seizure, or (4) acetaminophen + seizure. Imaging sessions were performed at scheduled intervals to minimize the effects of photobleaching and timed to capture periods before and after seizures to observe the potential effect of postictal hypoperfusion/hypoxia (Fig. 2). The imaging system consisted of a 2-photon microscope (Neurolabware) controlled by Scanbox (scanbox.org), a GUI that runs in Matlab. For calcium movie analysis, we used the SIMA python package, which includes motion correction and cell segmentation^81^. The signals were extracted, and spike probabilities were estimated using Scanbox^82^. Only cells that had spikes in the pre-seizure period were included in the analysis. Since some imaging sessions did not include periods of movement, we only analyzed data collected during immobility.

Blood vessel diameter was determined in ImageJ. The first 1000 frames of each session were used to generate a maximum intensity projection image. In other words, for a given pixel within a 1000 frame movie, the maximum pixel intensity at any time-point becomes that pixel’s intensity for the generated image. With this method, the neuropil and somas of the pyramidal cell layer appear bright, while blood vessels are dark. The mean change in blood vessel diameter for each imaging session was calculated from 3 identified blood vessels for control sessions and 5 identified blood vessels from seizure sessions.

To generate the plots of calcium activity during seizure and spreading depolarization, the extracted traces were converted to z-scored DF/F traces using a modified method of Jia et al 2011^83^. The time-dependent baseline was estimated for each cell by fitting a 3^rd^ order polynomial to the smoothed (100 frames symmetric sliding window average) F trace, excluding frames with running, negative peaks and the seizure period for the fit (frames 4000 to 6000 excluded from baseline fit for both seizure and control sessions). The z unit was determined by calculating the standard deviation (SD) of the DF/F trace, then recalculating the SD of the trace where it’s below 2 SD.

### 16-channel silicon probe LFP recordings from urethane-anaesthetized rats

16 adult male (300-400g) Long-Evans hooded rats (Charles River) were chronically implanted with an electrode for kindling in the right ventral hippocampus and an oxygen sensing probe in the right dorsal hippocampus as previously described^8^. The left side of the skull was marked for future acute recordings (3.5mm posterior, 2.2mm lateral to Bregma) by drilling a non-penetrating burr hole in the bone and marking with permanent marker. Thus, dental cement, jeweler screws, and a ground screw were confined to the right skull so as not to interfere with future microelectrode recordings. As with experiments performed in mice, rats recovered for 1-2 weeks and then underwent at least 5 once daily kindling sessions to confirm proper placement of the kindling electrode as evident by evoked seizures of stage 0-1 on the Racine scale and reliable oxygen recordings. The experiment began 2-3 weeks post-implantation.

On test day, rats were anesthetized with 1.2g/kg of urethane i.p. (Sigma) dissolved in saline and secured into a stereotaxic frame. As previously described^28^, a 16-channel linear microelectrode array with 100µm spacing (U-probe, Plexon Inc, Dallas, Texas) was slowly lowered into the hippocampus contralateral to the stimulation electrode and oxygen probe. Each contact was filtered between 0.1 and 500 Hz and amplified 1000X via a 16-channel headstage (unity gain) and amplifier (Plexon Inc.). All signals were referenced to ground and in turn, they were digitized at a sampling rate of 1kHz (Digidata 1440A + Axoscope; Molecular Devices, San Jose, CA). The probe was lowered until the apical dendritic border of the pyramidal cell layer (identified by extracellular reversal of theta rhythm) was situated in the dorsal third of the probe. Under anesthesia, 2 seconds of kindling stimulation at 1.5mA was used to elicit a seizure. Here, we analyzed data from a subset of rats that were injected with saline vehicle and experienced severe postictal hypoxia (n=4).

Raw LFP traces were visualized with AxoScope and signal analysis was conducted in Matlab (Mathworks, Natick, MA, USA)^28,29^, which was subsequently visualized using Origin (Microcal Software, Northampton, MA, USA). For the initial analysis, we selected the maximal amplitude signal from the probe, typically located close at the level of stratum lacunosum moleculare (SLM) or the hippocampal fissure. We computed spectrograms using a sliding window method using a 30s window separated by increments of 10s across the entire time series of the experiment. Individual spectra were computed within these 30s windows using a Welch’s averaged, modified periodogram method (pwelch) using 6s Hamming-windowed segments with 2s overlap. Spectra and spectrograms were inspected for characteristics of the activated (theta) or deactivated (SO) states by monitoring for peak logarithmic power in the 3 to 4.5 Hz versus 0.5 to 1.5 Hz bandwidths, respectively^28^. In most cases, regular temporal fluctuations of the log-transformed power values within the 0.5 to 1.5 Hz bandwidth of spectrograms were sufficient to characterize the state^29^. However, in some cases we used the ratio of log-transformed power values in the SO versus theta bandwidths since this tended to be a more sensitive index of state. Baseline and post-ictal samples of theta and/or SO activity were then used to characterize the properties and amplitudes of these state-dependent signals across time. In some cases, we could only analyze either one or the other state due to a lack of spontaneous samples occurring at an appropriate time point for analysis. For power measures, we extracted power at the relevant frequency peak for each state (theta 3.0 - 4.5Hz) and SO (0.5 - 1Hz) as well as in the gamma bandwidth (gamma during theta, 16.0 - 46.5Hz; gamma during SO, 14.0 - 39.5Hz). These data points were selected with respect to the timing of the hypoxic period. Maximal hypoxia occurred on average slightly less than 7 minutes following ictus and lasted up to an average of a full hour post-ictus. On average, measurements during maximal hypoxia were made 30 minutes post-ictus. The rising phase from maximal hypoxia started at an average of one hour following ictus and reached recovery phase after an average of approximately 100 minutes post-ictus. For the rising phase, measurements were made at an average of 80 minutes post-ictus while for the recovery phase, measurements were made at an average latency of 2 hours post-ictus.

For the same windowed segments of LFP, we used the full 16 traces from the probe to compute current source density (CSD), a spatial and temporal distribution of current sources and sinks underlying the voltage profile recorded^28,84^. The underlying assumptions for calculating CSD are based upon previous research^85-87^. To compute CSD, we made estimations of the second spatial derivative of voltage traces derived from the multiprobe based on a three-point difference (differentiation grid size of 300 µm) on the voltage values across spatially adjacent traces and expressed as mV/mm^2^, described previously in Wolansky et al^28^:

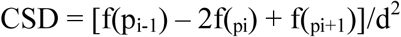

where f(pi) is the field potential signal from probe channel i (i = 2, 3, …, 15), and d is the distance between adjacent channels (0.1 mm). For traces at each end of the probe (i.e., channels 1 and 16), the differentiation grid was based solely on the immediately adjacent channel (e.g., channels 2 and 15, respectively). This latter procedure resulted in similar, if not identical, CSD results as the three-point differentiation method when we tested by successively eliminated probe end channels, re-computing and then comparing results obtained.

As with LFP spectral analysis, the CSD trace showing the largest fluctuations during both states (again at SLM) was spectrally analyzed throughout the entire experiment at the same time points pre and post-ictus as the LFP traces described above.

Cross frequency coupling of the maximal LFP signals was conducted using the method described by Tort et al^88^. Briefly, LFP traces during the baseline, hypoxic and recovery periods were zero-phase filtered in a 1Hz window across range of bandwidths from 0.5-10Hz at an interval of 0.25Hz to capture low frequencies. They were also filtered in a 5Hz window across a range of bandwidths from 10-200Hz at an interval of 2.5Hz to capture high frequencies. Via Hilbert transform, we extracted phase information from the low frequency signal and amplitude information from the fast frequency signal. The co-modulation of these respective low-to-high frequency signals was described via the modulation index (MI) and was plotted as a pseudo-colour 3-D plot to reveal any significant cross-frequency interactions. As previously described, the MI is a normalized index, with values ranging from 0 −1, and reflects the of the phase locking of fast frequency amplitude as a function of slow frequency phases^88^. We determined the frequencies of interest for each experiment to examine the co-modulation function in more detail. For theta, the low bandwidth was 2.5-5.5 Hz and the high bandwidth was between 20-45Hz. For SO, the low bandwidth was 0.1-2.5Hz and the high bandwidth was between 20-50Hz. By computing targeted phase-amplitude co-modulation plots between the low and high frequency Hilbert components for a minimum 40s signal during theta and 60s sample during SO, we were able to compare the phase preferences across our three conditions. These were assessed by computing the average preferred angle and radius for each normalized phase-amplitude co-modulation plot using the CircStat toolbox^89^. To test for any differences in the mean angle, we used the 95% confidence interval for the measure.

### LTP induction in urethane-anesthetized rats

Twenty-four Long-Evans rats (Charles River) previously chronically implanted with a bipolar stimulating electrode in the right ventral hippocampus and an oxygen probe in the dorsal right hippocampus (same coordinates as LFP experiment), were anesthetized with urethane (1.2 g/kg, i.p.) and positioned in a stereotaxic frame. All rats, including sham-treated, received 5-10 seizures prior to experimentation to ensure consistent seizure and hypoxia elicitation. All rats also participated in the object-context mismatch task prior to LTP experimentation, which takes two consecutive days. Body temperature was maintained by a heating pad with a temperature controller unit. Surgical procedure and field excitatory post-synaptic potential (fEPSP) recordings were performed in the left hemisphere of the brain as previously described^90^. Briefly, fEPSPs were evoked by stimulating the PP (AP:−7.9 L: 4.6 V: 2.6 - 3.2) with a stainless steel twisted bipolar electrode and recorded with an unipolar stainless-steel electrode implanted into the stratum lacunosum moleculare of the CA1 region of the hippocampus (AP: −3.2 L: 2.2 V: 2.6-2.8). During the surgical procedure, square-wave pulses of 0.2 ms duration were applied every 30 sec to the PP using a current stimulator (A-M system isolated pulse stimulator 2100). Both stimulating and recording electrodes were advanced slowly downward until reaching the optimal depth to record fEPSPs. After 60 min of post-surgical recovery, paired pulses were applied to record fEPSPs during the whole experiment which consisted of 10 min baseline, 30 min post-injection, 30 min post-seizure and 55 min post-HFS periods. Paired pulses were applied at different interpulse intervals (20, 25, 50, 100 ms) with a stimulus intensity set to evoke 40% of the fEPSP maximum slope.

Seizures were induced by delivering a kindling stimulation to the chronically-implanted electrode in the right hippocampus (2 s of 60 Hz biphasic 1 msec square wave pulses). HFS consisted of 10 trains of 15 pulses at 200 Hz, with 2 s delay between trains delivered to the PP using the same pulse parameters as in baseline. Signals were amplified using a MultiClamp 700B amplifier (Molecular Devices, high pass: 0.2 Hz, low pass: 5,000 Hz, gain: 200), digitized using an Axon Digidata 1440A data acquisition board (Molecular Devices) and stored on a personal computer using pClamp9 software (Molecular Devices). Sampling rate was set to 10 kHz.

### Object-context mismatch memory task

Twenty-four rats from the LTP experiment and an additional nineteen rats were used in this study. All rats were chronically implanted with oxygen probes and electrodes (same coordinates) and received 5-10 seizures before testing. Prior to testing, rats were pre-exposed to two different contexts devoid of any objects for 10 minutes each, one immediately after the other, each day for two consecutive days (habituation). Context A was a large white box (60cm x 60cm) housed in a well-lit room. Context B was a large black pentagonal-shaped bin (60cm in diameter) housed in a dimly-lit room. On the third day (test day), each context contained a unique pair of identical objects. The first object pair were small green ceramic cups, placed base down in the center of the arena. The second object pair were blue plastic pipette-tip holders, also placed in the center of the arena. On the testing day, a seizure was elicited (or sham) and 30 minutes following seizure termination rats were removed from the recording chamber and transported to the behavioural rooms. Rats investigated each context with paired objects for 5 minutes, one immediately after the other. Rats were then placed into their home cages for a 5-minute delay. Following this, rats were re-exposed to one of the contexts, this time with a single object from each pair placed in the center of the arena. Rats were given 3 minutes to explore, and an investigation ratio (time spent investigating the unfamiliar object/context pairing divided by total investigation time) was calculated. The context chambers, bedding, and objects were thoroughly wiped with 70% ethanol after each use.

## Acknowledgements

The authors thank Bonita Gunning and Sylwia Felong for their technical support. J.S.F. is supported by a Canadian Institutes for Health Research (CIHR) postdoctoral fellowship. R.C. is supported by an Eyes High postdoctoral fellowship funded by the University of Calgary. M.D.W. was funded by an Alberta Innovates graduate studentship. B.D. is funded by an American Epilepsy Society Postdoctoral Fellowship. Experiments in the laboratory of C.T.D., I.S., and G.C.T. were funded by Natural Sciences and Engineering Research Council of Canada (Discovery Grant #2016-06576), National Institutes of Health (Grant #NS99457), and CIHR (Grant # MOP-130495), respectively.

## Supplementary Material to

### Supplementary Figures

**Supplementary Figure 1:**
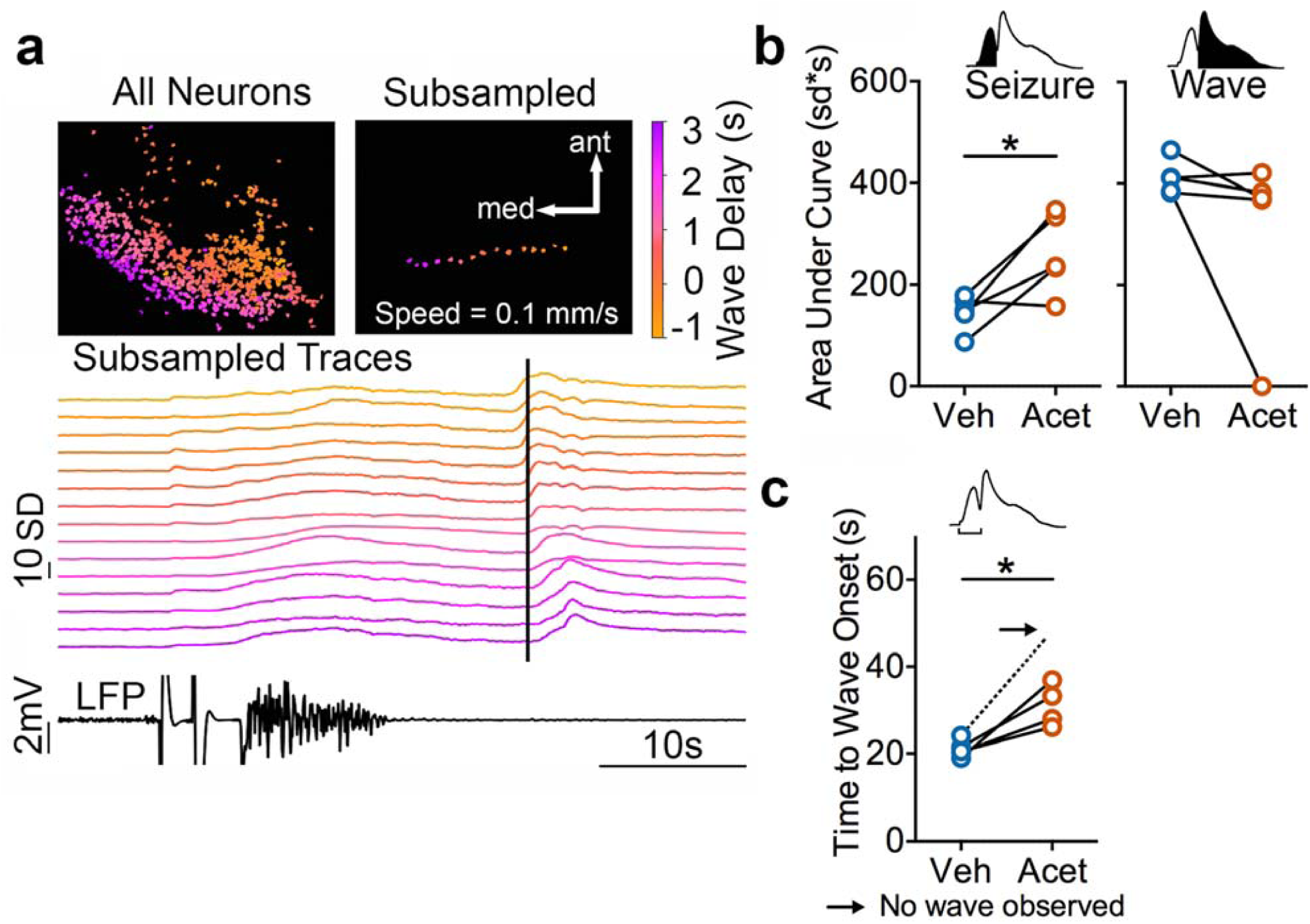
Seizure vs. wave dynamic alterations by acetaminophen. (**a**) Data from Figure 1c plotted to demonstrate spreading wave propagation direction and speed. For each recorded neuron, the color displayed corresponds to the delay of wave onset relative to the mean wave onset. Individual calcium traces from an unbiased subsampled group of 15 neurons show the direction of wave propagation travels towards the medial and posterior edges of the field of view. Arrows indicate medial and anterior directions. Vertical black bar indicates mean wave onset. (**b**) Quantification of calcium dynamics during seizures and calcium waves. Acetaminophen increased the area under the curve during the seizure (left; t(4)=3.14, p=0.03, paired t-test), but not the secondary wave (middle; t(4)=1.41, p=0.23). Y-axis applies to both graphs. (**c**) Acetaminophen significantly delayed the onset of the calcium wave appearance from seizure onset (paired t-test, t(3)=3.93, p=0.03. In one case (arrow, dotted line), no calcium wave appeared (data not included in analysis), which is also evident in the top and middle plots.

**Supplementary Figure 2:**
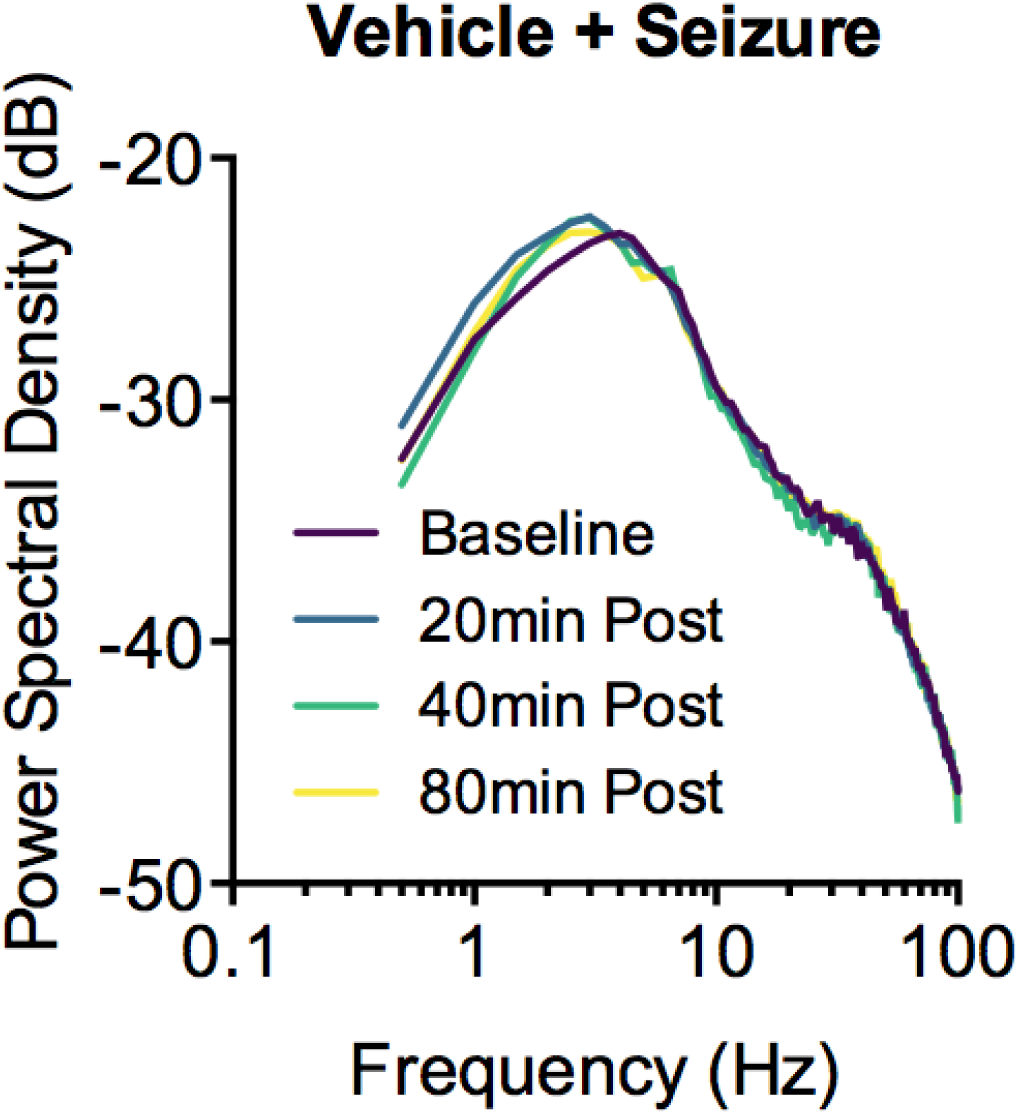
Postictal vasoconstriction does not alter the local field potentials in awake mice. Mean power spectral density plots are displayed before and 20, 40, or 80 minutes after a seizure. LFP was recorded as a differential recording between the two poles (tips 0.5 mm apart) of the simulating/recording electrode. Repeated measures ANOVA did not find an effect of time at the 0-4, 4-12, 25-55, and 55 100 Hz frequency bands.

**Supplementary Figure 3:**
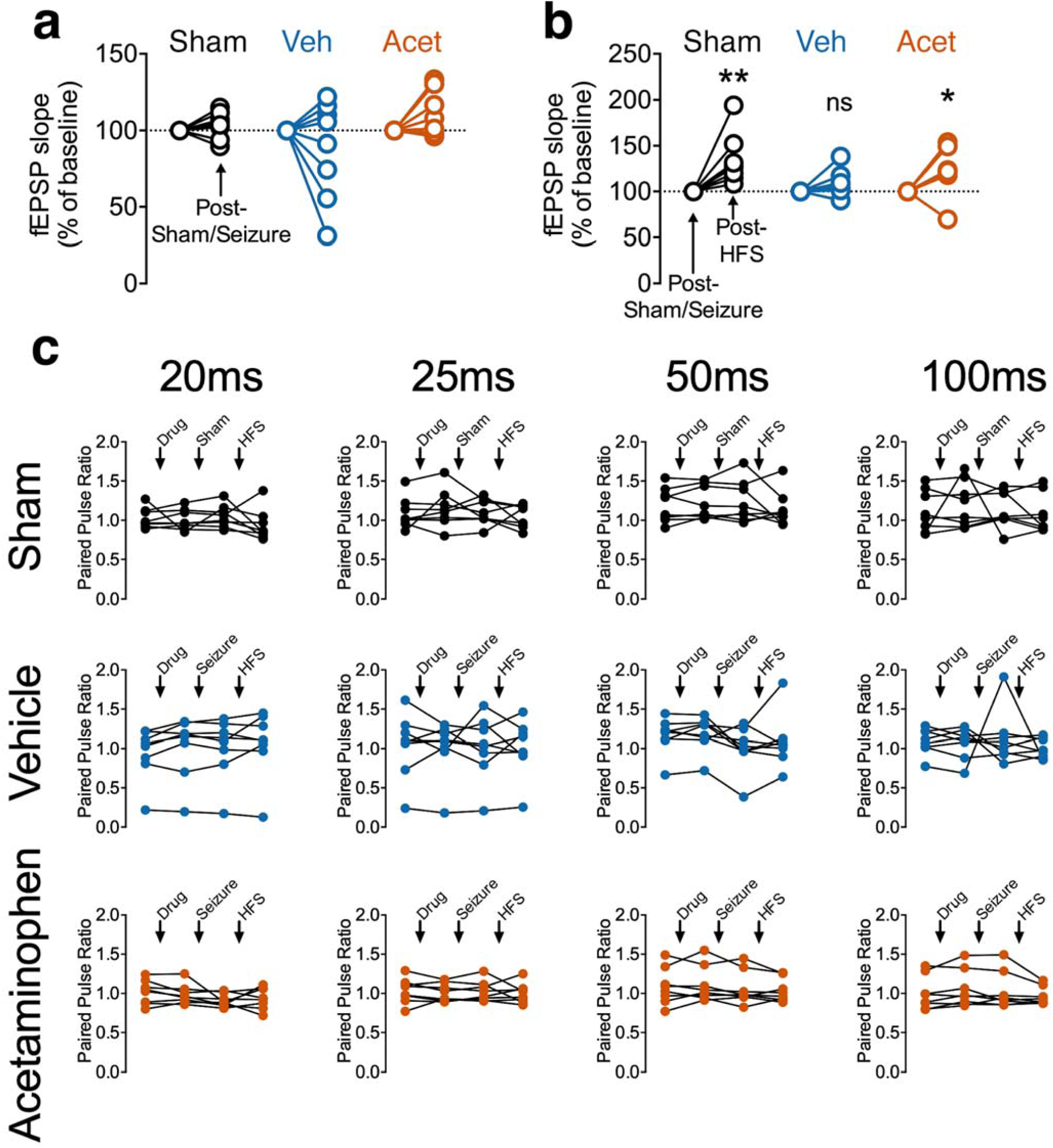
Seizures do not disrupt evoked responses or paired-pulse ratios. (**a**) No significant changes in fEPSP slope were observed following kindling stimulation (or sham) (sham in black - t(7)=1.21, p=0.27; vehicle in blue - t(7)=1.03, p=0.27; acetaminophen in orange - t(7)=1.96, p=0.09). (**b**) Data from Fig. 3c normalized to a baseline beginning after the seizure or sham. Significant potentiation was observed in sham controls (black - t(7)=3.60, p=0.0092) and rats who received acetaminophen prior to seizure induction (orange - t(7)=2.49, p=0.042), whereas no significant potentiation was observed in rats that received vehicle prior to seizure (blue - t(7)=1.81, p=0.11). (**c**) The paired pulse ratio from individual rats are plotted in a matrix according to the interval between the first and second pulse (20 −100ms) and treatment (Sham, Vehicle, or Acetaminophen). Paired one-way ANOVAs did not find a significant difference for any of the intervals or treatments.

**Supplementary Figure 4:**
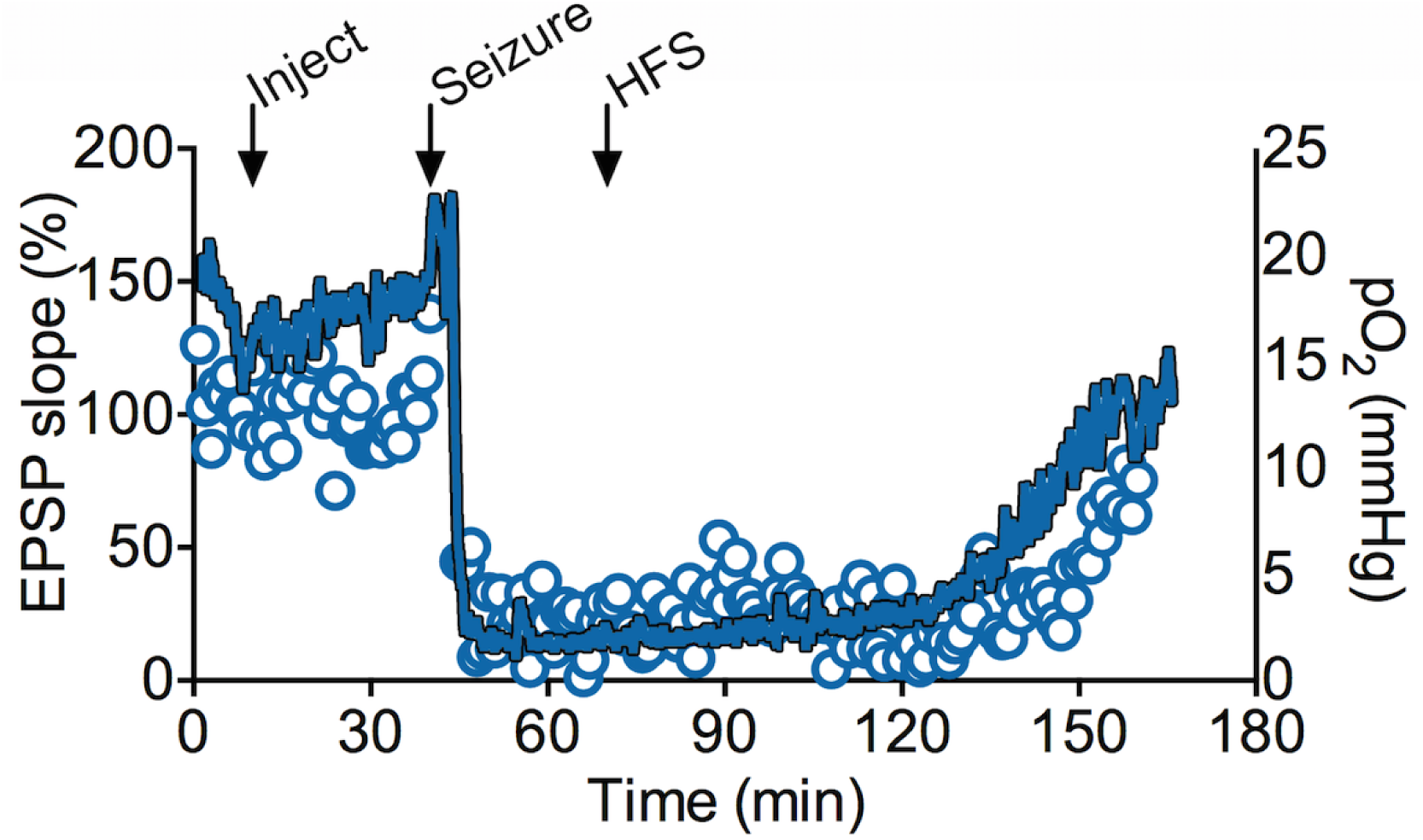
One exceptional example of suppression of evoked potentials following a seizure. An uncharacteristic example of a rat that experienced the most severe hypoxia in the experiment. The minimum pO_2_ reached 1.2mmHg and oxygen levels remained severely hypoxic (<10mmHg) for 102.8 minutes. Though LTP could not be induced, like other rats, a major reduction in evoked responses was uniquely noted during the entire postictal period. Note that the oxygen profile recovered before the evoked response recovered.

